# Biophysical Studies of Some Bee Products as Radioprotectors

**DOI:** 10.1101/472233

**Authors:** Shimaa F. Hamieda, Amal I. Hassan, Mona I. Abdou, Wafaa A. Khalil, Kamal N. Abd-El Nour

## Abstract

The study had been planned to evaluate some antioxidant ingredients in honey and propolis. Also, a study on ionizing gamma irradiated rats was done to assess these antioxidants as radioprotectors. Bioactive ingredients, such as phenols, flavonoids and trace elements, were explored using FTIR, UV-Vis and AAS spectroscopic techniques. Animals were exposed to fractionated gamma radiation doses. Honey, propolis and their combination were administrated before and during the irradiation period. Serum levels of total protein, albumin and uric acid were estimated. Also, the osmotic fragility of Red Blood Corpuscles (RBCs) membranes and a microscopic examination of blood films were investigated. The analysis demonstrated that the level of phenolic, flavonoid and trace elements are higher in propolis than honey. The levels of total protein and albumin decreased post irradiation while the level of uric acid increased. Likewise, osmotic fragility of RBCs membranes was increased with a sticking forming RBCs aggregation. It had been found that administration of the natural antioxidants induced amelioration in most of the studied parameters. It can be concluded that natural antioxidants produced a modulation against oxidative stress induced by ionizing radiation.

**Summary Statement:** Assessment of some antioxidant ingredients in honey and propolis. Also, a study on ionizing gamma irradiated rats was done to assess these antioxidants as radioprotectors.

## Introduction

Exposure to high doses of ionizing radiation is a rare event. Normally, occupational workers in different fields such as radiologists, industry, mining or research workers have restricted safety precautions. Although in some cases, such as radiation accidents high dose exposure may take place. Radio-protective agents have been widely examined to diminish the oxidative stress initiated by ionizing radiation (Saaya et al. 2017; Smith et al. 2017). Antioxidants are various types of molecules which in low levels fundamentally delays or prevents the oxidation effect of free radicals. (<u>Poljsak</u> et al. 2013). Reactive Oxygen Species (ROS) are produced from normal cellular metabolism (Khan et al. 2018) or as a consequence exposure to some chemicals and/or ionizing radiation (Hosseinimehr, 2010). Oxidative stresses can incentive to disturb redox balance between the production of ROS and the ability of cells to protect against them. Defense against oxidative stress kept up by utilizing several mechanisms which include antioxidant machinery (El-Missiry et al. 2007). The balanced cellular function is considered as a net result between the produced ROS and the available antioxidant defense mechanisms of the cell (Vit et al. 2010). Consequently, the decrease of cellular antioxidant capacity makes the biological system more susceptible to the malicious impacts of ROS (Birben et al. 2012). A great deal of research has been carried out on the radioprotective action of some synthetic chemical substances as antioxidants. These substances reduce mortality when administered pre exposure to lethal doses of ionizing radiation. But most of them have unwanted side effects that limited their use in medical practice (Cebolla et al. 2017). Using natural products as radioprotectors has several benefits since they are safe with proven therapeutic advantages. The body endogenous defensive system is supported by natural antioxidant compounds provided from nourishment (Nunes et al. 2013). Identification and isolation of antioxidants from natural sources has become an active field of research. Natural antioxidants can be micronutrients (vitamins and trace elements), phenolic compounds (flavonoids, and phenolic acids), nitrogen components (alkaloids, chlorophyll derivatives, amino acids, and amines), or carotenoids.

Various types of bees’ products, such as honey, pollen, royal gel, honey wax and propolis, have many therapeutic effects including anti-inflammatory, antimicrobial, antioxidant, antitumor, wound healing, and immunemodulatory activities (Boorn et al. 2010).

Honey is a sweet, viscous fluid, elaborated by bees from the nectar of plants and stored in their combs as food. Honey contains about 0.5% protein, mainly enzymes and amino acids (Khanal et al. 2010). Honey is readily available, affordable and well accepted by irradiated patients and useful for improving their lives (Orsolic et al. 2010). It is broadly accessible in many communities, although its mechanism of action remains unclear and requires more investigation.

Propolis is a resinous substance that bees collect from different plants. It is utilized in the construction of, and to seal the cracks in, the beehive. Chemical properties of propolis are not only advantageous to bees but have also pharmacological incentive as a natural mixture (Ali et al. 2010). It is a mixture of resin, basic oils and waxes (Cebolla et al. 2017). More than 200 constituents have been distinguished so far from propolis such as phenolic acids and their esters, caffeic acid and their esters, phenolic aldehydes, flavonoids and ketones; moreover, amino acids, proteins, vitamins (A, C, biotin, B_1_, B_2_ and B_3_), minerals (calcium, cobalt, copper, iron, magnesium, manganese, phosphorous, potassium, silicon and zinc) (Re et al. 1999). Phenolic compounds are able to scavenge ROS due to their electron donating properties. Also, their antioxidant adequacy relies on the dependability of various systems, besides the number and location of hydroxyl groups. Phenolic compounds exhibited higher antioxidant activity than carotenoids and vitamins (Moreira et al. 2011). Flavones are able to interact with free radicals and with the products of oxidative stress (Moreira et al. 2011; Treml and Smejkal, 2016). Flavonoids (including flavones, flavonols, flavanones and dihydroflavonols) and other phenolics (mainly substituted cinnamic acids and their esters) are the main active constituents of propolis and possess potent antioxidant activities (Almeida et al. 2011). Nutrient antioxidants may act together to reduce free radical’s levels (Stan et al. 2012).

Mineral elements are essential regulators of physiological processes. Calcium, zinc and magnesium are important as cofactors in enzymatic processes, mainly in the structure of the DNA repair system. Some minerals are components of important enzymes such as Zn for superoxide dismutase and Fe for catalase. Both enzymes protect the cell membranes from oxidative damage. Also Iron is a part of the heme of hemoglobin (Hb), myoglobin and cytochromes (Wołonciej et al. 2016). The high concentration of these mineral elements in propolis must promote the formation of these enzymes, and as a consequence provide its potent antioxidant capacity (Kocot et al. 2018).

The main goal of this work is to assess the level of some antioxidants constituents in honey and propolis such as phenols and trace elements using different spectroscopic techniques (*in vitro* study). Also it aimed to examine the extent of the healing ability of honey, propolis and their combination as radioprotectors against ionizing gamma radiation induced oxidative stress damages in male rats (*in vivo* study).

## Materials and methods

For **the in vitro studies**, three honey samples (H_1_, H_2_ and H_3_) and two propolis samples (P_1_ and P_2_) were used. H1 was from El-Fayoum area while the other two samples H2 and H3 were from the agricultural research center (Giza-Egypt). P_1_ was from agriculture research center - Giza-Egypt while P_2_ sample was from a local supermarket.

**UV-Visible (UV-Vis) spectroscopic techniques** type V-570 (Jasco, Germany) was used for the identification and quantification of phenolic compounds (Stan, 2012). Solutions of honey and propolis samples at a concentration of 0.1 mg/ml, and 10 mg/ml for honey only, were prepared as follows: 100 mg of honey and propolis are dissolved in 10 ml of distilled water and ethyl alcohol respectively in a ratio 1:1, then 1 ml of each solution diluted up to 100 ml of the same solvent. The absorbance between 200-600 nm was measured using the UV-visible spectrophotometer.

**Fourier Transform Infrared (FTIR) spectroscopic analysis** (Jasco FTIR 300 E, Japan) was chosen to explore the chemical composition of both honey and propolis samples for its straightforwardness and capacity to give fingerprints of the measured samples. Honey and propolis were dried at 150°C then grinded to fine powder, 2 mg of the powder sample was mixed with 200 mg KBr (FTIR grade) and pressed to make a pellet. The pellet was placed into the sample holder of FTIR to record the spectrum at the range of 4000-400 cm^−1^.

**Atomic absorption spectroscopy (AAS)** was used to study the mineral composition of honey and propolis. Six trace elements, Calcium (Ca), Sodium (Na), Iron (Fe), Magnesium (Mg), Potassium (K), and Zinc (Zn), were determined in three honey and two propolis samples. Samples were dried at 105°C and then mineralized by wet digestion method using (HNO_3_ - H_2_SO_4_). About 0.5 g of each sample was pre-digested in 4 mL of 65% HNO_3_ (Sigma-Aldrich, Germany) for 24 hours at room temperature, then 4 mL of 98% H_2_SO_4_ were added. After cooling, the solution was diluted to 20 mL with deionized water. Atomic absorption spectrophotometer (Agilent Technologies 200 series AA, USA) was used.

For **in vivo studies** Cobalt-60 Radiation source (gamma-cell 220), Atomic Energy of Canada Limited, installed at the Middle Eastern Regional Radioisotopes Center for the Arab Countries, Dokki, Cairo was used with an average exposure rate of 3.1 Gy per minute. Experimental rats were subjected to whole body irradiation with gamma fractionated doses of (1Gy / day for 5 continues days) i.e. 5 Gy total dose.

Eighty male albino rats weighing 150 −180 g from the National Research Center (Giza, Egypt) were used. The roles of the Medical Ethical Committee of the National Research Centre were taken in place. Animals were maintained for two weeks under acclimatization conditions of water and diet. They were divided into two main groups, control and irradiated ones. Each group was subdivided into four subgroups n=10 each. The four controls were the untreated one, the honey, propolis and their combination treated subgroups. The four irradiated subgroups were the irradiated one, and the subgroups which irradiated and protected with honey, propolis and their combination.

**Honey** was diluted with water and administered orally to animals at a dose of 250 mg/kg/day in a volume of 1 ml/rat for 15 continuous days of the honey sub-control groups. The irradiated subgroups received honey 10 days before irradiation and 5 days during the fractionated irradiation doses.

**Propolis** was extracted using 70% ethanol; about 10 g of propolis was dissolved in 100 ml ethanol. The supernatant was separated using filter paper Whatman No (1). The extract was completely evaporated under reduced pressure. Propolis was freshly prepared pre-oral administration in saline at a dose of 90 mg/kg/day for 15 continuous days of the sub-propolis control group. The irradiated subgroups received propolis 10 days before irradiation and 5 days during the fractionated irradiation doses.

**Honey and propolis combined subgroups** received 250 mg/kg honey plus 90 mg/kg propolis /day for continuous 15 days of the combined control sub-group. The irradiated subgroup received the same dose for 10 days before irradiation and 5 days during the fractionated irradiation doses.

**Blood samples** were collected from orbital venous plexuses of the rat eye at different time intervals 1^st^, 7^th^ and 14^th^ days post last dose of irradiation. Serum samples were stored at −20°C for the biochemical investigations. Heparinized blood samples were collected for osmotic fragility and blood films were stained for microscopic examination.

**Biochemical analysis** of total protein was determined according to Gornal et al. (1949). Albumin was determined according to Gendler (1984). Uric acid was determined according to Bahram and Trinder (1972). Commercial kits from Biodiagnostic Company, Egypt were used.

**The osmotic fragility of Red Blood Corpuscles test** is used to detect the fragility of RBCs of different groups. Whole blood was added to varying percent buffered sodium chloride solution of concentrations 0.900, 0.765, 0.675, 0.585, 0.540, 0.500, 0.450, 0.400, 0.360, 0.315, 0.270, 0.180, 0.090 and 0.000 and allowed to incubate at room temperature. The amount of hemolysis is then determined by reading the absorbance of the supernatants at 540 nm on the spectrophotometer Unicam UV-Visible Spectrometry (Helios, United Kingdom). Normal and treated blood samples had been tested at the same condition (Brown, 1993).

The percent of hemolysis of different samples was calculated for each supernatant as follow:

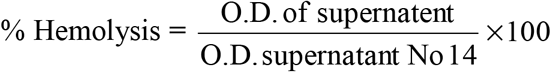

O.D. is the Optical Density of supernatant of different NaCl concentrations. O.D. supernatant No 14 is the O.D. of 0.000 NaCl concentrations which represents 100% hemolysis.

**Blood films** were stained with Leishman stain for 5 min then washed with water for examination. They were scanned and examined using the light microscope (PB362040x) at a magnification power 400 x. The images were captured using a digital camera (Yashica, EZ8032, 8.2 megapixels).

**Statistical analysis** of the biochemical data was performed by two-way analysis of variance (ANOVA) test and Duncan test using SPSS software program version 17 (SPSS Inc, USA).

## Results

**UV - Visible spectroscopic analysis** results are appeared in figure (1A & 1B). Figure (1A) represents a simple spectrophotometric registration of UV-visible spectra of different honey and propolis samples at a concentration of 0.1 mg / ml. The first propolis sample spectrum has an absorption band with ***λ*_max_** at 288 nm. The second propolis sample spectrum has a plateau with ***λ*_max_** between 270-290 nm. Both propolis samples have an absorption peak at 230, while honey samples did not show any absorption band in this region. At a concentration of 10 mg / ml, honey samples showed spectra with ***λ*_max_** around 280 nm, while the absorption peak at 230 nm only observed in H1 sample as appeared in Figure (1B).

**Figure (1):**
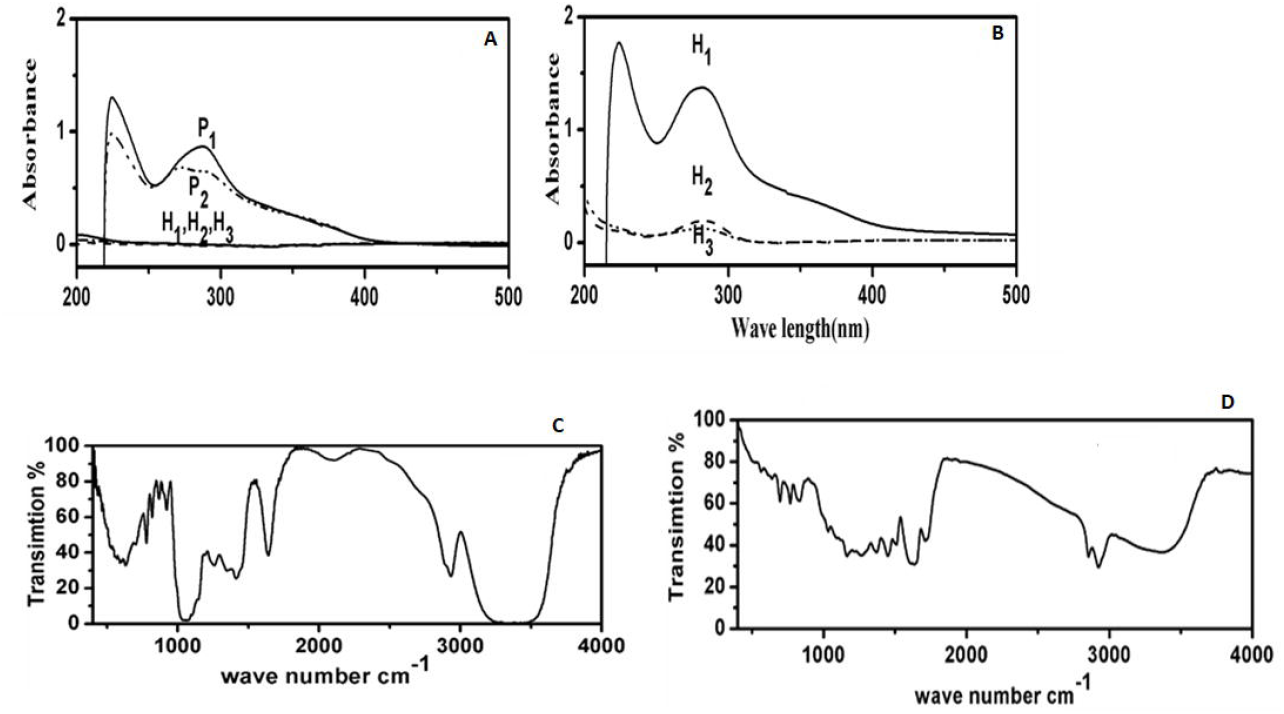
Absorption spectra of flavonoid (A) UV-Vis of different honey and propolis samples at concentration 0.1 mg / ml and (B) UV-Vis of different honey samples at concentration 10 mg /ml (C) FTIR spectrum pattern of honey, (D) FTIR spectrum pattern of propolis.

**FTIR spectroscopic analysis** shows the spectra pattern of honey in Figure (1C) and of propolis in Figure (1D). The presence of distinct bands in the spectra pattern is considered to be an indication of the presence of certain functional group and the expected compounds are listed in Table (1). The variations in the intensity of such bands of different samples were listed in Table (2). The correlation coefficient (R^2^) of FTIR band intensity is represented in Table (3).

**Table (1):**
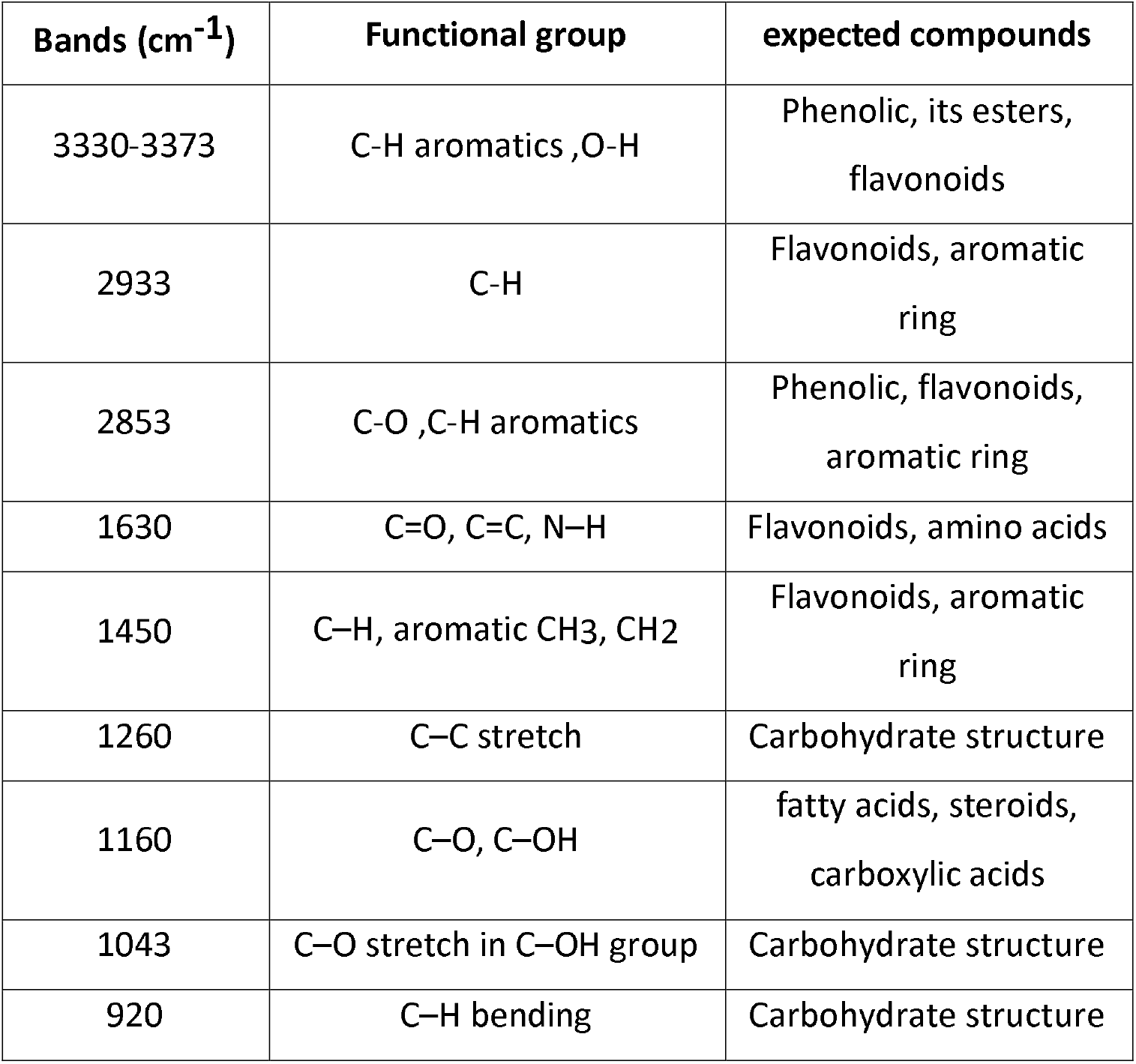
Distinct bands of honey and propolis in FTIR spectra

**Table (2):** The variation of bands intensity in FTIR spectra of honey and propolis

| Sample<br>Band (cm<sup>-1</sup>) | H2 | H3 | P1 | P2 |
| --- | --- | --- | --- | --- |
|  | Intensity |  |  |  |
| 3373 | 0.67 | 1.56 | 44.45 | 36.60 |
| 2933 | 27.55 | 26.80 | 41.65 | 29.45 |
| 2853 | — | — | 48.38 | 34.49 |
| 1630 | 39.69 | 38.47 | 43.58 | 30.85 |
| 1450 | 30.28 | 25.29 | 49.37 | 34.78 |
| 1260 | 34.15 | 32.72 | 49.74 | 35.01 |
| 1160 | — | — | 47.55 | 34.76 |
| 1043 | 4.16 | 1.70 | 55.20 | 46.66 |

**Table (3):**
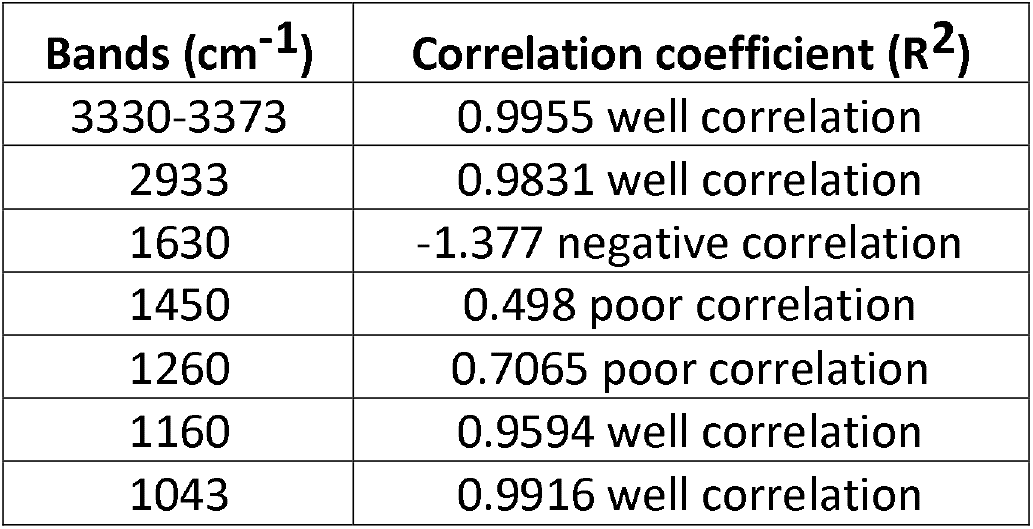
C orrelation coefficient of FTIR bands intensity

| Bands (cm<sup>-1</sup>) | Correlation coefficient (R<sup>2</sup>) |
| --- | --- |
| 3330-3373 | 0.9955 well correlation |
| 2933 | 0.9831 well correlation |
| 1630 | -1.377 negative correlation |
| 1450 | 0.498 poor correlation |
| 1260 | 0.7065 poor correlation |
| 1160 | 0.9594 well correlation |
| 1043 | 0.9916 well correlation |

The total flavonoid and phenols in P_1_ sample is found to be slightly higher than that P_2_. H1 sample contains higher content of flavonoid and phenols than H2 and H3 samples. Also, it has been found that propolis has greater contents of flavonoid and phenols than honey.

### Mineral composition

Honey and propolis mineral composition was measured using atomic absorption spectroscopy. Six elements (Fe, Mg, Na, K, Ca and Zn) were determined in three honey and two propolis samples. The concentrations of these minerals in honey and propolis samples are listed in Table (4). One-way ANOVA test demonstrated a significant difference at p < 0.05 in the content of each mineral amongst propolis and honey samples as well as between different honey or propolis samples.

**Table (4):** Concentration of the trace elements studied in honeys and propolis in mg/100g dry weight.

| Mineral | Sample |  |  |  |  |
| --- | --- | --- | --- | --- | --- |
|  | H1 | H2 | H3 | P1 | P2 |
| Fe | 1.97±0.3<sup>d</sup> | 1.51±0.8<sup>e</sup> | 3.58±0.6<sup>c</sup> | 37.98±2.4<sup>a</sup> | 28.78±1.2<sup>b</sup> |
| Mg | 7.78±1.6<sup>c</sup> | 4.04±2.9<sup>c</sup> | 7.57±2.4<sup>c</sup> | 153.24±9.1<sup>a</sup> | 41.60±1.7<sup>b</sup> |
| Na | 3.60±0.4<sup>b</sup> | 1.89±0.4<sup>b</sup> | 3.77±0.7<sup>b</sup> | 18.80±1.9<sup>a</sup> | 17.96±1.1<sup>a</sup> |
| K | 17.63±0.4<sup>c</sup> | 14.5±0.8<sup>cd</sup> | 9.15±1.6<sup>d</sup> | 105.82±7.4<sup>a</sup> | 75.37±2.4<sup>b</sup> |
| **Ca** | 35.56±1.9<sup>cd</sup> | 39.10±1.7<sup>c</sup> | 27.65±1.4<sup>d</sup> | 155.30±6.6<sup>a</sup> | 76.54±1.2<sup>b</sup> |
| **Zn** | 0.62±0.2<sup>c</sup> | 0.11±0.1<sup>c</sup> | 0.13±0.1<sup>c</sup> | 24.63±1.2<sup>a</sup> | 13.43±0.5<sup>b</sup> |

**Data expressed as mean ± SD, mg/100g of dried sample, n = 3**
**\* Significant different at p < 0.05.** Duncan test results are expressed as letters (a, b, c, d and e).
Values with different superscript letters are significantly differ. Values with same letters indicate no significant difference between samples. For each element separately (each row).

### Biochemical analysis

The data of serum total protein and albumin were summarized in Tables (5) and (6). A two-way ANOVA test and Duncan test indicated a significant decrease in their levels (P<0.05) in irradiated rats contrasted to the normal group. Also, treatments with honey, propolis, and their combination induced significant ameliorations compared to irradiated rats. ANOVA showed no significant change between different time intervals post irradiation (P> 0.05) of total protein and albumin and also no significant interaction between factors (P> 0.05).

**Table (5):** Total serum protein levels in irradiated rats and those treated with honey, propolis separately and combined pre *γ*- irradiation.

| Treatment | Days after γ- irradiation | | |
| --- | --- | --- | --- |
|  | 1<sup>st</sup> | 7<sup>th</sup> | 14<sup>th</sup> |
| **Normal** | 5.44±0.26<sup>a</sup> | 5.50±0.87<sup>a</sup> | 5.34±0.85<sup>a</sup> |
| **Honey** | 6.00±0.30<sup>a</sup> | 5.44±0.14<sup>a</sup> | 4.65±0.71<sup>a</sup> |
| **Propolis** | 4.81±0.74<sup>ab</sup> | 5.30±0.23<sup>ab</sup> | 4.60±0.68<sup>ab</sup> |
| **Combined** | 5.00±0.16<sup>ab</sup> | 5.23±0.17<sup>ab</sup> | 5.04±0.34<sup>ab</sup> |
| **Irradiated** | 3.78±0.34<sup>c</sup> | 3.06±0.49<sup>c</sup> | 3.34±0.31<sup>c</sup> |
| **Irradiated + honey** | 4.97±0.78<sup>b</sup> | 4.30±0.55<sup>b</sup> | 4.21±0.81<sup>b</sup> |
| **Irradiated + prpolis** | 5.23±0.68<sup>ab</sup> | 5.49±0.63<sup>ab</sup> | 5.00±0.45<sup>ab</sup> |
| **Irradiated + combined** | 5.44±0.53<sup>a</sup> | 5.37±0.35<sup>a</sup> | 5.24±0.82<sup>a</sup> |

**Table (6):** Serum albumin levels in irradiated rats and treated with honey, propolis separately and combined pre *γ*- irradiation.

| Treatment | Days after γ- irradiation | | |
| --- | --- | --- | --- |
|  | 1<sup>st</sup> | 7<sup>th</sup> | 14<sup>th</sup> |
| Normal | 3.1±0.102<sup>a</sup> | 2.88±0.44<sup>a</sup> | 2.94±0.44<sup>a</sup> |
| Honey | 3.42±0.21<sup>a</sup> | 3.17±0.08<sup>a</sup> | 2.66±0.42<sup>a</sup> |
| Propolis | 3.23±0.39<sup>a</sup> | 3.00±0.08<sup>a</sup> | 3.21±.42<sup>a</sup> |
| Combined | 3.2±0.44<sup>a</sup> | 3.1±0.39<sup>a</sup> | 3.2±0.197<sup>a</sup> |
| Irradiated | 2.54±0.13<sup>b</sup> | 1.96±0.04<sup>b</sup> | 1.68±0.12<sup>b</sup> |
| Irradiated + honey | 3±0.10<sup>a</sup> | 2.93±0.44<sup>a</sup> | 2.95±0.46<sup>a</sup> |
| Irradiated + prpolis | 3.19±0.11<sup>a</sup> | 2.90±0.40<sup>a</sup> | 3.50±0.11<sup>a</sup> |
| Irradiated + combined | 2.82±0.43<sup>a</sup> | 2.99±0.11<sup>a</sup> | 2.95±0.43<sup>a</sup> |

The data of serum uric acid are summarized in Tables (7). A two-way ANOVA test and Duncan test indicated that uric acid significantly increased (P<0.05) in rats exposed to γ-radiation compared with the normal group. Also, different treatments with honey, propolis, and their combination induced significant amelioration compared to irradiated group. The propolis and combination pretreated animal groups showed improvement in serum uric acid levels more than the honey pretreated group. Also, ANOVA demonstrated no significant difference between the level of uric acid at various time intervals post irradiation (P> 0.05) and furthermore no significant interaction between factors (P> 0.05).

**Table (7):** Serum uric acid levels in irradiated rats and those treated with honey, propolis separately and combined pre *γ*- irradiation

| Treatment | Days after γ- irradiation | | |
| --- | --- | --- | --- |
|  | 1<sup>st</sup> | 7<sup>th</sup> | 14<sup>th</sup> |
| **Normal** | 1.43±0.04<sup>b</sup> | 1.32±0.03<sup>b</sup> | 1.48±0.08<sup>b</sup> |
| **Honey** | 1.38±0.03<sup>b</sup> | 1.51±0.031<sup>b</sup> | 1.59±0.05<sup>b</sup> |
| **Propolis** | 1.46±0.09<sup>b</sup> | 1.56±0.03<sup>b</sup> | 1.38±0.11<sup>b</sup> |
| **Combined** | 1.51±0.11<sup>b</sup> | 1.43±0.05<sup>b</sup> | 1.45±0.08<sup>b</sup> |
| **Irradiated** | 2.28±0.03<sup>a</sup> | 3.00±0.05<sup>a</sup> | 3.23±0.19<sup>a</sup> |
| **Irradiated +**<br>**honey** | 1.41±0.09<sup>b</sup> | 1.40±0.07<sup>b</sup> | 1.52±0.06<sup>b</sup> |
| **Irradiated +**<br>**prpolis** | 1.48±0.06<sup>b</sup> | 1.38±0.04<sup>b</sup> | 1.38±0.05<sup>b</sup> |
| **Irradiated +**<br>**combined** | 1.38±0.05<sup>b</sup> | 1.17±0.04<sup>b</sup> | 1.15±0.08<sup>b</sup> |

## Biophysical examinations

### Red blood cells membranes studies

#### Osmotic fragility

Figures (2A-H) shows the effect of NaCl percent concentrations on osmotic fragility of RBCs membrane and its differentiation.

**Figure (2):**
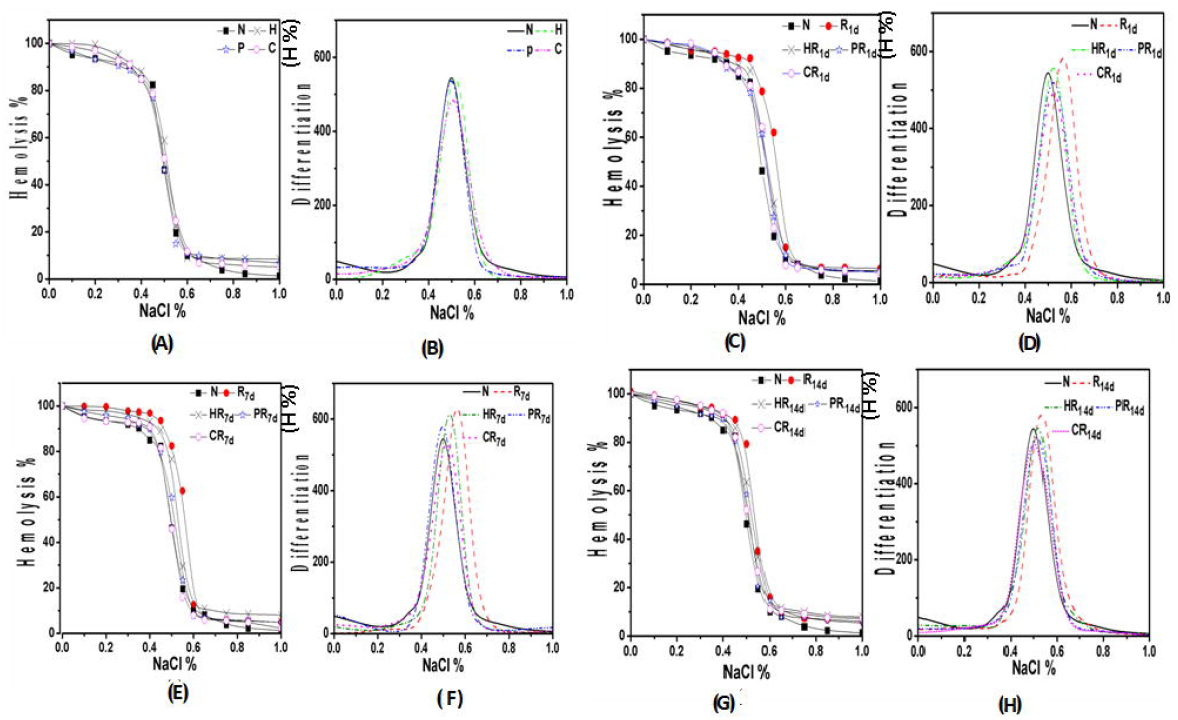
(2 A and B) RBCs of control (A) Osmotic fragility curves and (B) Differentiation curves. (2 C and D) RBCs of 1^st^ day post-irradiation (C) Osmotic fragility curves and (D) Differentiation curves (E and F) RBCs of 7^th^ day post- irradiation (E) Osmotic fragility curves and (F) Differentiation curves (2 G and H) RBCs of 14^th^ day post- irradiation (G) Osmotic fragility curves and (H) Differentiation curves.

Measurements illustrate the variation of the percentage hemolysis as a function of the percentage of NaCl concentration in buffer solution for RBCs of different animal subgroups as appeared in Figures (2 A, C, E and G).

To analyze these data, the graphs were differentiated and plotted as a function of the percentage of NaCl concentration as shown in Figures (2 B, D, F and H). From osmotic fragility curves, it is possible to calculate Median Corpuscular Fragility (MCF) which means the NaCl concentration at which 50% of RBCs are hemolyzed. The points of the differentiation curves were calculated by subtracting each point from the previous one in the hemolysis curve. The width at half maximum (W**_hmax_**) of the differentiation curve represents the relative elastic limit of RBCs membrane. The increase of W**_hmax_** expresses more elasticity of cell membrane. The MCF and W**_hmax_** values of RBCs from different subgroups were represented in Table (8).

**Table (8):** The values of MCF and Wh_max_ for from different animal subgroups RBCs

| **Group** | **MCF** | **Whmax(arbitrary units)** |
| --- | --- | --- |
| **Control group** |  |  |
| **N** | 0.498 | 0.140 |
| **H** | 0.514 | 0.140 |
| **P** | 0.498 | 0.139 |
| **C** | 0.500 | 0.163 |
| **1<sup>st</sup> day post irradiation** |  |  |
| **R1d** | 0.570 | 0.135 |
| **HR1d** | 0.520 | 0.139 |
| **PR1d** | 0.520 | 0.149 |
| **CR1d** | 0.520 | 0.150 |
| **7<sup>th</sup> days post irradiation** |  |  |
| **R7d** | 0.560 | 0.130 |
| **HR7d** | 0.525 | 0.130 |
| **PR7d** | 0.515 | 0.140 |
| **CR7d** | 0.498 | 0.150 |
| **14<sup>th</sup> days post irradiation** |  |  |
| **R14d** | 0.530 | 0.132 |
| **HR14d** | 0.520 | 0.137 |
| **PR14d** | 0.515 | 0.147 |
| **CR14d** | 0.500 | 0.150 |

### Blood film

Histological examinations of RBCs by light microscope are shown in Figures (3 A-P). The images of the control samples appear in Figures (3 A-D), for negative control and positive control administered honey propolis and their combination respectively, have normal structures. Figures (3 E-H, and 3 M-P) illustrate blood film images of the irradiated and irradiated pretreated animal subgroups at different time intervals 1^st^, 7^th^ and 14^th^ day post-irradiation respectively. The results showed a remarkable sticking of RBCs forming aggregation, which persist until the 14^th^ day post-irradiation Figures (3 E, I and M). Oral administration of propolis and the combination diminishes the formation of RBCs aggregation and reduces the damage of irradiation Figures (3 G, K and O) and (3 H, I and P). Instead, honey recovers RBCs at the 14^th^ day post irradiation Figure (3 N) while at 1^st^ and 7^th^ days Figures (3 F and J) still have some sort of sticking and deformation.

**Figure (3 A-P):**
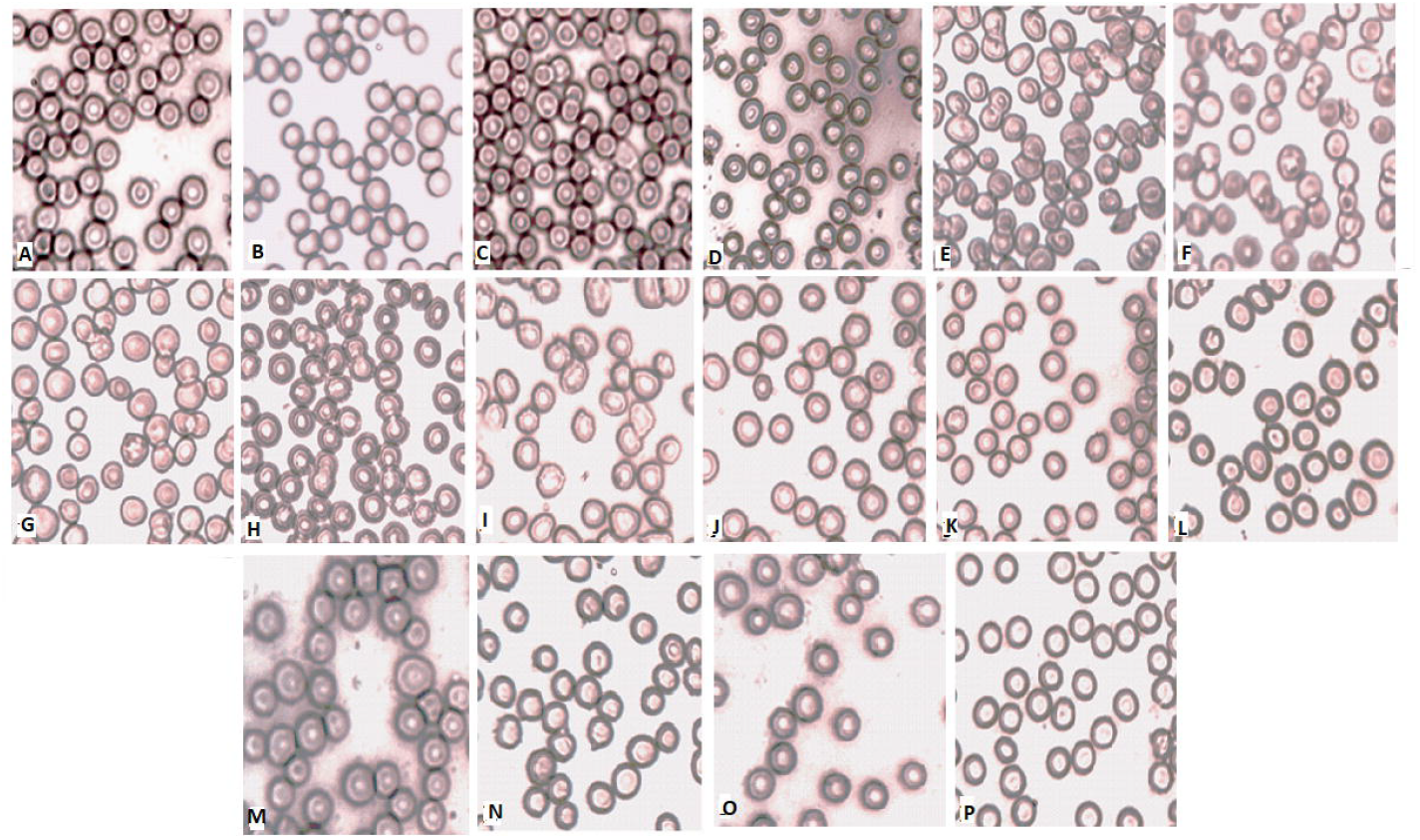
Microscopic blood film observations of different animal groups RBCs (400X) and (96DPI). (3 A-D) Microscopic blood film observations of controls (A) Negative control rats, (B) Control rats administered honey, (C) Control rats administered propolis (D) Control rats administered combination. (3 E-H) Microscopic blood film observations of 1^st^ -day post irradiation animal group; (E) irradiated (f) irradiated + honey (G) irradiated + propolis (H) irradiated + combination. (3 I-L) Microscopic blood film observations of 7^th^ day post irradiation animal group; (I) irradiated (J) irradiated + honey (K) irradiated + propolis (L) irradiated + combination. (3 M-P) Microscopic blood film observations of 14^th^ day post irradiation animal group; (M) irradiated (N) irradiated + honey (O) irradiated + propolis (P) irradiated + combination.

## Discussion

UV - Vis spectroscopic investigation affirmed the presence of phenolic compounds in honey and propolis since they exhibit two major absorption bands in UV - Vis region (Moţ et al., 2011). The characteristic bands of phenols are a broad band centered around 280–330 nm and a second band at 250 nm (Yang, al., 2013).

The UV-Vis spectrum in Figure (1a) is comparable with an earlier research work which isolates and evaluates six antioxidants of propolis at (200-600 nm) (Yang, al., 2013). The spectra of propolis at a concentration of 0.1 mg/ml are similar in shape and position to the spectra of honey at a concentration of 10 mg/ml Figure (1b). These results are in coincidence with beforehand published works (Mihai, et al., 2011).

In general, it is thought that the appearance of the absorption band at 270-330 nm is attributable to flavonoids and phenols (Mărghitaş et al., 2013) while the absorption band at 230 nm is characteristic for flavone/flavonol by-products (Yang, t al., 2013). Also, it is reported that band with λ_max_ 290 nm, is believed to arise from Hispertin (Makawi et al., 2009). Variation in intensity and shift in the absorption band might be attributed to the different antioxidant ratios in flavonoids and phenols (Mărghitaş et al., 2013). A high correlation was found between the UV-Vis spectra of samples and total phenolic compounds that were determined by the Folin-Ciocalteu method (Hamieda et al., 2015).

FTIR spectroscopic data in Tables (1) and (2) are comparable with those early reported (Gallardo-Velázquez et al., 2009; Ali, et al., 2012). The bands in Table (2) showed different correlations to antioxidant activity represented in Table (3) which is in coincidence with Mot and his coworkers’ results (Moţ et al., 2011). It is clear that the antioxidant activity of polyphenols does not depend only on the amount of the present compounds but also on their chemical structures (Omene et al., 2012).

The Fe, Mg, Na, K, Ca and Zn are basic elements since they play important roles in the biological systems one of them is their antioxidant activity (Pohl et al., 2012).

The higher minerals concentration in propolis than honey is in coincidence with previously reported data (Grembecka and Szefer 2013; Formicki et al., 2013). Also, honey and propolis samples differ in their mineral constituents depending on the resources in the soil and kind of plants from which the bees took nectar (Shah et al., 2014). These elements do many biological functions since they act as coenzymes and participate in other processes like Redox one (Kurek-Górecka et al., 2013).

The biochemical analysis of serum total protein is relatively important in assessing the health state of mammals (Vasile et al., 2009). The decrease of total protein post irradiation might be due to the disturbances of the vital biological processes or because of progress in the penetrability of the liver, kidney and other tissues resulting in leakage of protein via the kidney (Muhammad et al., 2015). The decline in the level of albumin concentration could be due to enhanced degradation as well as enhanced loss of albumin through the gastrointestinal tract (Shabon, 2005). Moreover, the extensive damage to the lymphoid organs following *γ*-irradiation was probably related to the drop in serum globulin levels (Moulder et al., 2004). Earlier investigations by Wheeler and Bernard (1994) have reported that irradiation leads to proteinuria, which is associated with low serum total protein and albumin. An improvement in the levels of serum total proteins and albumin in the pretreated groups with propolis and combination maintain their levels near the normal level at the different time intervals. The pretreatment with honey only shows slight improvement. The effect of pretreatment with propolis is in harmony with Saleh (2012) who reported that propolis significantly improved the total protein content of the liver and kidney and indicated more profound therapeutic effects.

The increased uric acid level in rats exposed to γ-radiation compared with the normal group is supported by other works (Ferreira-Leach and Hill, 2001; Saleh, 2012). Uric acid is the end product of the purine metabolism and is normally eliminated by the kidney in urine. Excess uric acid may be a side effect of some cancer treatments and may lead to a condition called tumor lysis syndrome (Crohns et al., 2009). The excess uric acid forms crystals which may deposit in the tiny tubes of the kidney and cause acute kidney damage, leading ultimately to kidney failure (Abd elhalim and Moussa, 2013). This result might be ascribed to the hindrance of glomerular selective properties caused by irradiation (Berry et al., 2001).

The mechanism by which the natural product honey and propolis anticipate renal oxidative stress may incorporate several mechanisms. One of them is the induction of glutathione GSH synthesis which is consumed when bind to free radicals. Others are by a scavenger effect of antioxidants phenols and flavonoids (naringenin, pinostrombin and galangin) found in honey and propolis to the electrons of ROS. This effect could have accumulated in the cells of the proximal convoluted tubules of the kidney where propolis was reported to be collected and secreted (Saleh et al., 2013). Instead of ROS, their free electrons will be captured with the antioxidant (Newairy, et al., 2009). So there must be a massive concern to clarify the clinical role of the propolis.

Osmotic fragility of RBCs membranes showed an increase in hemolysis percentage and osmotic fragility of RBCs membrane in irradiated groups which was detected through the shift of the osmotic fragility curves towards the higher NaCl concentration. Also, the reduction in W**_hmax_** is an evidence of decreasing the flexibility of RBCs. This result is in coincidence with Selim and his coworkers (Selim et al., 2009). It is interesting to confirm that these changes in osmotic fragility can be omitted by pretreating with honey, propolis, and their combination.

The increase in the osmotic fragility may be attributed to some changes in the properties of the RBCs membrane. Ionizing radiation causes a disturbance in energy metabolism, disorganization of lipoprotein structure of biological membranes, peroxidation of membrane lipids and inactivation of various bound enzymes (Na^+^, K^+^, and Mg^+^) ATPase. It has also been suggested that radiation induces perturbation in Na^+^ and K^+^ transport system which is connected with oxidation of membrane protein sulfhydryl groups (Jóźwiak and Helszer 1981). Modification in the physical condition of the membrane proteins may lead to change in the permeability of the RBCs membrane. Some proteins of the cell membrane act as pores (ion channels) through which the aqueous solutions carried inside the cell (Hemida et al., 2011).

In the presence of honey, propolis and their combination protective effect against RBCs membrane damage were observed. This observation could be attributed to the presence of some coenzyme minerals such as Zn for superoxide dismutase and Fe for catalase. Both enzymes protect the cell membrane from oxidative damage. Since propolis contains both elements, it promotes the formation of the dismutase and catalase and as a consequence decreases the hemolysis. Additionally, Potassium is the principle cation in intracellular fluid and functions in the regulation of the osmotic pressure and co-enzyme of Na^+^/K^+^ ATPase (Achikanu et al., 2013; Shah, et al., 2014). It was found that protective effect of propolis on RBCs membrane was related to the higher phenolic content of propolis (Moreira et al., 2011; Valente et al., 2011). These results mean that propolis act as a powerful antioxidant compound which prevents the oxidative damage of lipoproteins of the cell membrane (Yousif et al., 2012).

RBCs are well known that they are large cells having a biconcave structure which is naturally flexible and bendable (El-marakby et al., 2013). They do not stick together as a result of the Coulomb repulsive forces between the positive electrostatic charges on their outer surface membranes. The electrostatic charges are formed on the surface of the normal healthy RBCs subsequently of the K ion pump which forms the resting potential across cellular membranes (Hemida et al., 2012). RBCs cell membranes are particularly sensitive to the effects of irradiation. The sticking of several cells together after irradiation is due to the oxidative damage of the free radicals. They destruct the processes within both layers of RBCs membrane and that alters the membrane structure and function (El-marakby et al., 2013). Also, the sticking can be attributed to the changes in the packing properties of the phospholipid bilayer and macromolecules forming the cellular membrane. This will cause changes in the membrane permeability to ion transport and the liquid crystalline phase of the membrane will change. Therefore, one may state that changes in the membrane permeability will result in the change of the membrane bioelectric potential and surface electrostatic charges (Hemida et al., 2012). The loss or decrease of the surface electrostatic charges on the cellular membrane will deteriorate the repulsive forces between adjacent RBCs membranes and cause the sticking. Moreover, the oxidation of membrane proteins leads to red cell vesiculation (Moreira et al., 2011). It is noticed that pretreating of the animals with honey decreases the sticking of cells and reaching mostly to normal shape by propolis and their combination. This is because the hydroxyl groups of the phenolic and flavonoids compounds have anti-lipid peroxidant and rheologically protecting against oxidative stress (de Kok et al., 2010).

## Conclusion

The administration of natural antioxidants such as honey and propolis mixture mitigates γ- induced oxidative stress in rat blood.

UV absorption spectra and dielectric measurement were found to be good tools to support the data given by biochemical analysis, such as total proteins. The obtained data indicated that honey and propolis mixture display an in vivo antioxidant activity, which is evidenced experimentally by ameliorating the osmotic fragility, sticking and aggregation of red blood cell as well as serum total protein and albumin.

Further investigations are required to elucidate the mechanisms of propolis and honey actions.

## References

Abd elhalim, M. A. K., & Moussa, S. A. A. (2013). The biochemical changes in rats’ blood serum levels exposed to different gamma radiation doses. African Journal of Pharmacy and Pharmacology, 7(15), 785–792.

Achikanu, C. E., Eze-Steven, P. E., Ude, C. M., & Ugwuokolie, O. C. (2013). Determination of the vitamin and mineral composition of common leafy vegetables in south eastern Nigeria. Int. J. Curr. Microbiol. App. Sci. 2(11), 347–353.

Ali, F.H., Kassem, G. M. and Atta-Alla, O. A. (2010). Propolis as a natural decontaminant and antioxidant in fresh oriental sausage, Vet. Ital. 46, 167–172.

Ali, I. H., Daoud, A. S., & Shareef, A. Y. (2012). Physical properties and chemical analysis of Iraqi propolis. Tikrit Journal of Pure Science, 17(2), 26–31.

Almeida, I. M., Barreira, Oliveira, J. C., & Ferreira, I. C. (2011). Dietary antioxidant supplements: benefits of their combined use. Food and Chemical Toxicology, 49(12), 3232–3237.

Berry, C., Brosnan, M. J., Fennell, J., Hamilton, C. A., & Dominiczak, A. F. (2001). Oxidative stress and vascular damage in hypertension. Current opinion in nephrology and hypertension, 10(2), 247–255.

Bhram, D. and Trinder, P. (1972). An improved colour reagent for the determination of blood glucose by the oxidase system. Analyst,1972, 97, 1425.

Birben, E., Sahiner, U. M., Sackesen, C., Erzurum, S. and Kalayci, O. (2012). Oxidative Stress and Antioxidant Defense. World Allergy Organ J. 5(1), 9–19.

Boorn, K., Khor, Y., Sweetman, E., Tan, F., Heard, T. and Hammer, K. (2010). Antimicrobial activity of honey from the stingless bee Trigona carbonaria determined by agar diffusion, agar dilution, broth microdilution and time-kill methodology. J. Appl. Microbiol. 108, 1534–1543.

Brown BA. Hematology: Principles and procedures. 6th edn. Philadelphia, USA: Lea and Febiger, 1993.

Cebolla, A., Demarzo, M., Martins, P., Soler, J., Garcia-Campayo, J. (2017). Unwanted effects: Is there a negative side of meditation? A multicentre survey. PLoS ONE, 12(9), e0183137.

Crohns, M., Saarelainen, S., Kankaanranta, H., Moilanen, E., Alho, H., & Kellokumpu-Lehtinen, P. (2009). Local and systemic oxidant/antioxidant status before and during lung cancer radiotherapy. Free radical research, 43(7), 646–657.

de Kok, T. M., de Waard, P., Wilms, L. C., & van Breda, S. G. (2010). Antioxidative and antigenotoxic properties of vegetables and dietary phytochemicals: the value of genomics biomarkers in molecular epidemiology. Molecular nutrition & food research, 54(2), 208–217. Eferewr

El-marakby S. M., N. S. Selim, O. S. Desouky, H. A. Ashry, A. M. Sallam, (2013). Effects of Poly-MVA on the rheological properties of blood after in-vivo exposure to gamma radiation, Journal of radiation research and applied sciences, (6), 2, 21–30.

El-Missiry, M. A., Fayed, T. A., El-Sawy, M. R., & El-Sayed, A. A. (2007). Ameliorative effect of melatonin against gamma-irradiation-induced oxidative stress and tissue injury. Ecotoxicology and Environmental Safety, 66(2), 278–286.

Ferreira-Leach, A. M. R., & Hill, E. M. (2001). Bioconcentration and distribution of 4-tert-octylphenol residues in tissues of the rainbow trout (Oncorhynchus mykiss). Marine environmental research, 51(1), 75–89.

Formicki, G., Gren, A., Stawaez, R., Zysk, B., & Gal, A. (2013). Metal content in honey, propolis, wax, and bee Pollen and implications for metal pollution monitoring. Pol. J. Environ. Stud. 22, 99–106.

Gallardo-Velázquez, T., Osorio-Revilla, G., Zuñiga-de Loa, M., & Rivera-Espinoza, Y. (2009). Application of FTIR-HATR spectroscopy and multivariate analysis to the quantification of adulterants in Mexican honeys. Food Research International, 42(3), 313–318.

Glender, S. (1984). Proteins. In: Clinical Chemistry: Theory, Analysis and Correlation, Kaplan, L.A. and A.J. Pesce (Eds.). Mosby CV. Elsevier, Toranto. 1268–1327.

Gornal, A. C., Bardawill, C. J. and David, M. M. (1949). Determination of serum proteins by means of the biuret reaction. J. Boil. Chem. 177, 751–766.

Grembecka, M., & Szefer, P. (2013). Evaluation of honeys and bee products quality based on their mineral composition using multivariate techniques. Environmental monitoring and assessment, 185(5), 4033–4047.

Hamieda, S. F., Hassan, A. I., Abdou, M. I., Khalil, W. A., & Abd-El nour, K. N. (2015). Evaluation of radioprotective effects of some bee products and its flavonoid constituents: in vivo study on male rats. Rom. J. Biopys. 25(1), 13–34.

Hemida, S. F., Abd-El-Nour, K. N., Farag, H., Aiad, T. H. A., Alrouby, M., & Hassan, N. S. (2012). Role of melatonin in reducing morphological changes in red blood cell induced by gamma irradiation. Rom. J. Biopys. 22(3-4), 221–234.

Hemida, S., El nour, K. N. A. B. D., Farag, H., Aiad, T. A., & Alrouby, M. (2011). Protective effect of melatonin on hemoglobin damage induced by gamma irradiation. Rom. J. Biophys. 21(4), 317–329.

Hosseinimehr, S. J. (2010). Flavonoids and genomic instability induced by ionizing radiation. Drug Discovery Today, 15(21), 907–918.

Jóźwiak, Z., & Helszer, Z. (1981). Participation of free oxygen radicals in damage of porcine erythrocytes. Radiation Res. 88(1), 11–19.

Khan, F., Garg, V. K., Singh, A. K., Kumar, T. (2018). Role of free radicals and certain antioxidants in the management of huntington’s disease: a review. J. Analytical & Pharmaceutical Res. 7(4), 386–392.

Khanal, B., Baliga, M. and Uppal, N. (2010). Effect of topical honey on limitation of radiation-induced oral mucositis: an intervention study, Int. J. Oral Maxillofac. Surg. 2010, 39, 1181–1185.

Kocot, J., Kiełczykowska, M., Luchowska-Kocot, D., Kurzepa, J. and Musik, I. (2018). Antioxidant Potential of Propolis, Bee Pollen, and Royal Jelly: Possible Medical Application. Oxid. Med. Cell. Longev. 2018, 7074209.

Makawi, S. Z. A., Gadkariem, E. A., & Ayoub, S. M. H. (2009). Determination of antioxidant flavonoids in Sudanese honey samples by solid phase extraction and high performance liquid chromatography. Journal of Chemistry, 6(S1), S429–S437.

Mărghitaş, L. A., Dezmirean, D. S., & Bobiş, O. (2013). Important developments in Romanian propolis research. Evidence-Based Complementary and Alternative Medicine, 2013.

Mihai, C. M., Mărghitaş, L. A., Dezmirean, D. S., & Bărnuţiu, L. (2011). Correlation between polyphenolic profile and antioxidant activity of propolis from Transylvania. Scientific Papers Animal Science and Biotechnologies, 44(2), 100–103.

Moreira, L. L., Dias, T., Dias, L. G., Rogão, M., Da Silva, J. P., & Estevinho, L. M. (2011). Propolis influence on erythrocyte membrane disorder (hereditary spherocytosis): A first approach. Food and Chemical Toxicology, 49(2), 520–526.

Moreira, L.L., Dias, T., Dias, L. G., Rogao, M., Silva, J. P., Estevinho, L. M. (2011). Propolis influence on erythrocyte membrane disorder (hereditary spherocytosis): A first approach. Food Chem. Toxicol. 49, 520–526.

Moţ, A. C., Silaghi-Dumitrescu, R., & Sârbu, C. (2011). Rapid and effective evaluation of the antioxidant capacity of propolis extracts using DPPH bleaching kinetic profiles, FT-IR and UV–VIS spectroscopic data. Journal of Food Composition and Analysis, 24(4), 516–522.

Moulder, J. E., Fish, B. L., & Cohen, E. P. (2004). Impact of angiotensin II type 2 receptor blockade on experimental radiation nephropathy. Radiation Res. 161(3), 312–317.

Muhammad, M. M. A., Mouchira, M., & Naglaa, R. A. (2015). Physiological effects of Bee Venom and Propolis on irradiated Albino rats. Danish Journal of Agriculture and Animal Sciences, 11–21.

Newairy, A. S. A., Salama, A. F., Hussien, H. M., & Yousef, M. I. (2009). Propolis alleviates aluminum-induced lipid peroxidation and biochemical parameters in male rats. Food and Chemical Toxicology, 47(6), 1093–1098.

Nunes, L. C., Galindo, A. B., Lustosa, S. R., Brasileiro, M. T., Do Egito, A. A., Freitas, R. M., Randau, K.P. Rolim Neto, P. J. (2013). Influence of seasonal variation on antioxidant and total phenol activity of red propolis extracts. Adv. Studies Biol. 5, 119–133.

Omene, C. O., Wu, J., & Frenkel, K. (2012). Caffeic Acid Phenethyl Ester (CAPE) derived from propolis, a honeybee product, inhibits growth of breast cancer stem cells. Investigational New Drugs, 30(4), 1279–1288.

Orsolic, N. (2010). A review of propolis antitumor action in vivo and in vitro, J. Api Prod. Api. Med. Sci. 2, 1–20.

Pohl, P., Stecka, H., Sergiel, I., & Jamroz, P. (2012). Different aspects of the elemental analysis of honey by flame atomic absorption and emission spectrometry: a review. Food Analytical Methods, 5(4), 737–751.

Poljsak, B., Šuput, D. and Milisav, I. (2013). Achieving the Balance between ROS and Antioxidants: When to Use the Synthetic Antioxidants. Oxid Med Cell Longev. 2013, 956792.

Re, R., Pellegrini, N., Proteggente, A., Pannala, A., M., Yang, M. and Evans, C. R. (1999). Antioxidant activity applying an improved ABTS radical cation decolorization assay, Free Radic. Biol. Med. 26, 1231–1237.

Saaya, F. M., Katsube, T., Yi Xie, Y., Tanaka, K., Fujita, F. and Wang, B. (2017). Research and Development of Radioprotective Agents: A Mini-Review. International Journal of Radiology 4(2-3), 128–138.

Saleh, E. M. (2013). Antioxidant effect of aqueous extract of propolis on hepatotoxicity induced by octylphenol in male rats. Acta Toxicol. Argent, 20(2), 68–81.

Saleh, O. M., Soliman, M. M., Mansour, A. A. K., & Abdel-Hamid, O. M. (2013). Protective effects of propolis on gamma-irradiated nigella sativa extract induced blood and immune changes in wistar rats. American Journal of Biochemistry & Biotechnology, 9(2), 162.

Selim, N. S., Desouky, O. S., Ali, S. M., Ibrahim, I. H., & Ashry, H. A. (2009). Effect of gamma radiation on some biophysical properties of red blood cell membrane. Romanian J Biophys, 19, 171–185.

Shabon, M. H. (2005). Radioprotective effects of soya and garlic oils in irradiated male rats. Isotope Radiation Research, 37, 1525–1534.

Shah, A., Sikandar, F., Ullah, I., Khan, S. U. D., Rana, U. A., & McCoy, T. (2014). Spectrophotometric Determination of Trace Elements in various Honey Samples, Collected from different Environments. Journal of Food and Nutrition Research, 2(9), 532–538.

Smith, T. A., Kirkpatrick, D. R., Smith, S., Smith, T. K., Pearson, T., Kailasam, A., Herrmann, K. Z., Schubert, J. and Agrawal, D. K. (2008). The effects of Yucca schidigera and Quillaja saponaria on DNA damage, protein oxidation, lipid peroxidation, and some biochemical parameters in streptozotocin-induced diabetic rats. Journal of Diabetes and its Complications, 22(5), 348–356.

Stan, L. (2012). Propolis commercial tinctures–phenolics and anti-oxidant activity. Agricult. Pract. Sci. J. 83,(3-4).

Treml, J. and Smejkal, K. (2016). Flavonoids as Potent Scavengers of Hydroxyl Radicals. Comprehensive Reviews in Food Science and Food Safety 15, 720–738.

Valente, M. J., Baltazar, A. F., Henrique, R., Estevinho, L., & Carvalho, M. (2011). Biological activities of Portuguese propolis: protection against free radical-induced erythrocyte damage and inhibition of human renal cancer cell growth in vitro. Food and Chemical Toxicology, 49(1), 86–92.

Vasile, M., Teren, O., Ciupina, V., & Turcu, G. (2009). Changes of electrophoretical fractions in simultaneous exposure to gamma radiation and hyperbarism. Romanian Reports in Physics, 61(1), 121–128.

Vit, P., A. Rodriguez-Malaver, C. Rondon, I. Gonzalez, M. Luisa, DI Bernardo, M. Ysabal Garcia, (2010). Bioactive indicators related to bioelements of eight unifloral honeys, Arch. Latinoam. Nutr. 60, 405–410.7

Wheeler, D. C., & Bernard, D. B. (1994). Lipid abnormalities in the nephrotic syndrome: causes, consequences, and treatment. Am. J. Kidney dis. 23(3), 331–346.

Wołonciej, M., Milewska, E., Roszkowska-Jakimiec, W. (2016). Trace elements as an activator of antioxidant enzymes. Postepy Hig. Med. Dosw. 70(0),1483–1498.

Yang, L., Yan, Q.H., Ma, J.Y., Wang, Q., Zhang, J.W., Xi, G.X. (2013). High performance liquid chromatographic determination of phenolic compounds in propolis. Tropical Journal of Pharmaceutical Res. 12(5): 771–776.

Yousif Y, B., Sanaa, S., Talal, A., & Shtywy, A. (2012). Structure-activity relationships regarding the antioxidant effects of the flavonoids on human erythrocytes. Natural Sci. 2012.

Yousif Y, B., Sanaa, S., Talal, A., & Shtywy, A. (2012). Structure-activity relationships regarding the antioxidant effects of the flavonoids on human erythrocytes. Natural Sc. (4) 9, 740–747.

